# Myosin VI and β-arrestin synergistically regulate GIPR internalization and signaling

**DOI:** 10.1101/2025.08.22.671816

**Authors:** Nishaben M. Patel, Sivaraj Sivaramakrishnan

## Abstract

Glucose-dependent insulinotropic peptide receptor (GIPR) stimulates insulin release and regulates metabolic homeostasis. GIPR function is shaped by spatiotemporal trafficking of this G protein-coupled receptor (GPCR). While GPCR endocytosis is traditionally associated with β-arrestin, GIPR internalization is only modestly dependent on this pathway. In this study, we demonstrate that GIPR engages a cytoskeletal motor, myosin VI to drive receptor endocytosis. GIPR engages the adaptor-motor complex through a PDZ-binding motif (PBM) at its C-ail. Interestingly, β-arrestin binding to phosphorylated residues upstream of the PBM enhance myosin VI recruitment and activation. GIPR internalization is dependent on both receptor phosphorylation and the PBM site to recruit β-arrestin and myosin VI, respectively. Synergistic engagement of β-arrestin and myosin VI results in desensitization of GIP-stimulated cAMP signaling while activating pERK1/2 from endosomal compartments. Blocking myosin VI activity enhances insulin release in pancreatic beta cells, demonstrating a novel role for this pathway in regulating the physiological effects of GIPR. Our findings highlight the direct convergence of two independent trafficking pathways at the level of the receptor C-tail, with implications for the nuanced regulation of individual GPCRs through the differential engagement of β-arrestin and myosin VI.

**Significance:** GIPR has emerged as a frontline drug target in type 2 diabetes and obesity. Cellular effects of GIPR are regulated by receptor internalization and desensitization through mechanisms that are unclear. Here, we identify a novel GIPR trafficking pathway through the engagement of a cytoskeletal motor, myosin VI. Myosin VI and β-arrestin synergistically regulate GIPR endocytosis, signaling and insulin response in pancreatic beta cells. Our study highlights the convergence of two parallel trafficking mechanisms in GPCR function with potential implications in targeting metabolic disorders.

## Introduction

G protein-coupled receptor (GPCR) internalization is an established mechanism for receptor desensitization, compartmentalized signaling, and spatiotemporal regulation of physiological responses (1, 2). Clathrin-dependent GPCR endocytosis typically proceeds through the recruitment of β-arrestin to a barcode of GRK phosphorylated residues in the GPCR C-tail (3). In parallel, ∼ 30% of GPCR C-tails contain a PDZ-binding motif (PBM) that can recruit PDZ adaptor proteins to facilitate receptor trafficking (4–7). The relative effects of β-arrestin and PDZ adaptors on GPCR internalization is dependent on the specific sequence motifs in the receptor C-tail (8, 9). The PBM is typically located in the last 3-5 amino acids of the receptor C-tail (8). In contrast, GRK phosphorylation sites are dispersed throughout the C-tail and their differential phosphorylation mediates distinct downstream signaling effects through conformational effects on β-arrestin (9). While these two pathways have been extensively studied, their potential intersection in regulating signaling and physiological responses in the same receptor is unclear and forms the premise of this study.

This study focuses on glucose-dependent insulinotropic peptide receptor (GIPR), whose differential desensitization is of interest in targeting metabolic disorders (10). GIPR is a class B GPCR that regulates glucose metabolism, body weight, bone formation and food intake (11, 12). GIPR, along with its counterpart GLP1R, is responsible for the increase in insulin secretion from beta cells in the islets of Langerhans postprandially (13). This process is referred to as the incretin effect and is compromised in type 2 diabetes (T2D) (14). Incretin receptors are prime targets of weight loss and T2D medications (12). Incretin mimetics have transformed the medical landscape for treatment of obesity and T2D. Both agonism and antagonism at GIPR reported positive effects on body weight (10). Chronic agonism of GIPR desensitizes cAMP signaling response of the receptor (15). Additionally, co-agonism at GIPR and GLP1R has achieved superior weight loss results, compared to mono-agonism at GLP1R (12, 16). These contrasting results highlight the gap in our knowledge on GIPR function and regulation and further insight into GIPR trafficking is required to reconcile these previous findings.

Membrane trafficking of GIPR is an essential factor in shaping the signaling and physiological response of this receptor (17). GIPR is constitutively internalized and recycled through the fast-recycling pathway (18). Binding of GIP reroutes the receptor to the trans Golgi network (TGN) and the slow-recycling pathway (18). The molecular mechanisms that govern GIPR trafficking are relatively unexplored. Additionally, the pathways that govern differential signaling of therapeutic drugs, compared to the native agonist, are unknown. Further insights into the regulatory pathways of GIPR that guide receptor trafficking and recycling, desensitization, and drug tolerance are required to delineate mechanisms of action of medications for obesity and T2D.

GIPR has an extensive C-tail with multiple phosphorylation sites that recruit β-arrestin and a putative PDZ binding motif (PBM) (8, 19). We have demonstrated previously that GPCRs can recruit myosin VI through the PBM site (20). Therefore, in this study, we use GIPR to delineate the intersection between two trafficking pathways - β-arrestin and myosin VI. We demonstrate that β-arrestin and myosin VI activity are both required for GIPR internalization, cAMP production and phosphorylation of ERK. β-arrestin enhances binding of GIPC, an adaptor of myosin VI, to the phosphomimetic GIPR C-tail, stimulating motor activity. Finally, we show that myosin VI-dependent internalization desensitizes GIPR-mediated insulin secretion in pancreatic beta cells. Together, our data support a model wherein GIPR internalization is synergistically regulated by β-arrestin and myosin VI, leading to desensitization of insulin secretion.

## Results

### GIPR internalization requires both β-arrestin and myosin VI

- GIP stimulation of GIPR has been shown to enhance receptor internalization in a β-arrestin2 dependent manner (18). GIPR has a type III PDZ binding motif (X-X-C) that we have recently shown to engage a cytoskeletal trafficking pathway in the D2 dopamine receptor (D2R) (8, 20). To examine the relative roles of these two mechanisms, we used a myosin VI-selective inhibitor (TIP) and a β-arrestin dual-knockdown cell line (Δβ-arr) (Fig. 1A) (21, 22). GIP treatment (1 μM; 15 min) stimulated GIPR internalization, as visualized using antibody-labeled GIPR in HEK293 cells (Fig. 1B). Consistent with previous studies, constitutive knockdown of β-arrestin (Δβ-arr) yields reduced internalization (Fig. 1B), as quantified from the number of internalized puncta per cell (Fig. 1C) and cytosol-to-surface ratio of fluorescence intensity (Fig. 1D) (23). TIP pretreatment (100 μM; 15 min) also abolishes GIPR internalization (Fig. 1B-D) with no further effects observed with both Δβ-arr and TIP treatment. These data demonstrate that both β-arrestin and myosin VI activity are necessary for GIP-stimulated GIPR internalization. GIP-stimulated internalization has been shown to result in reduced cAMP signaling from the remaining cell surface receptors (24). Accordingly, blocking GIPR internalization through TIP pretreatment (10 μM; 15 min) in HEK293 cells with transient GIPR overexpression, significantly enhances cAMP accumulation in response to GIP treatment (1 μM; 15 min) (Fig. 1E). Interestingly, these augmented cAMP levels with TIP pretreatment are observed both in WT and Δβ-arr cells (Fig. 1E). GIP stimulation is known to trigger ERK1/2 phosphorylation with a peak response at 4 min (25). We observe pERK1/2 stimulation in HEK293 cells with transient GIPR overexpression both in WT and Δβ-arr cells (Fig. 1F). In both cell types, TIP pretreatment significantly diminishes pERK1/2 levels. Given the lack of internalization upon TIP treatment (Fig. 1B-D), we infer that MAPK activity stems from internalized receptor.

**Figure 1:**
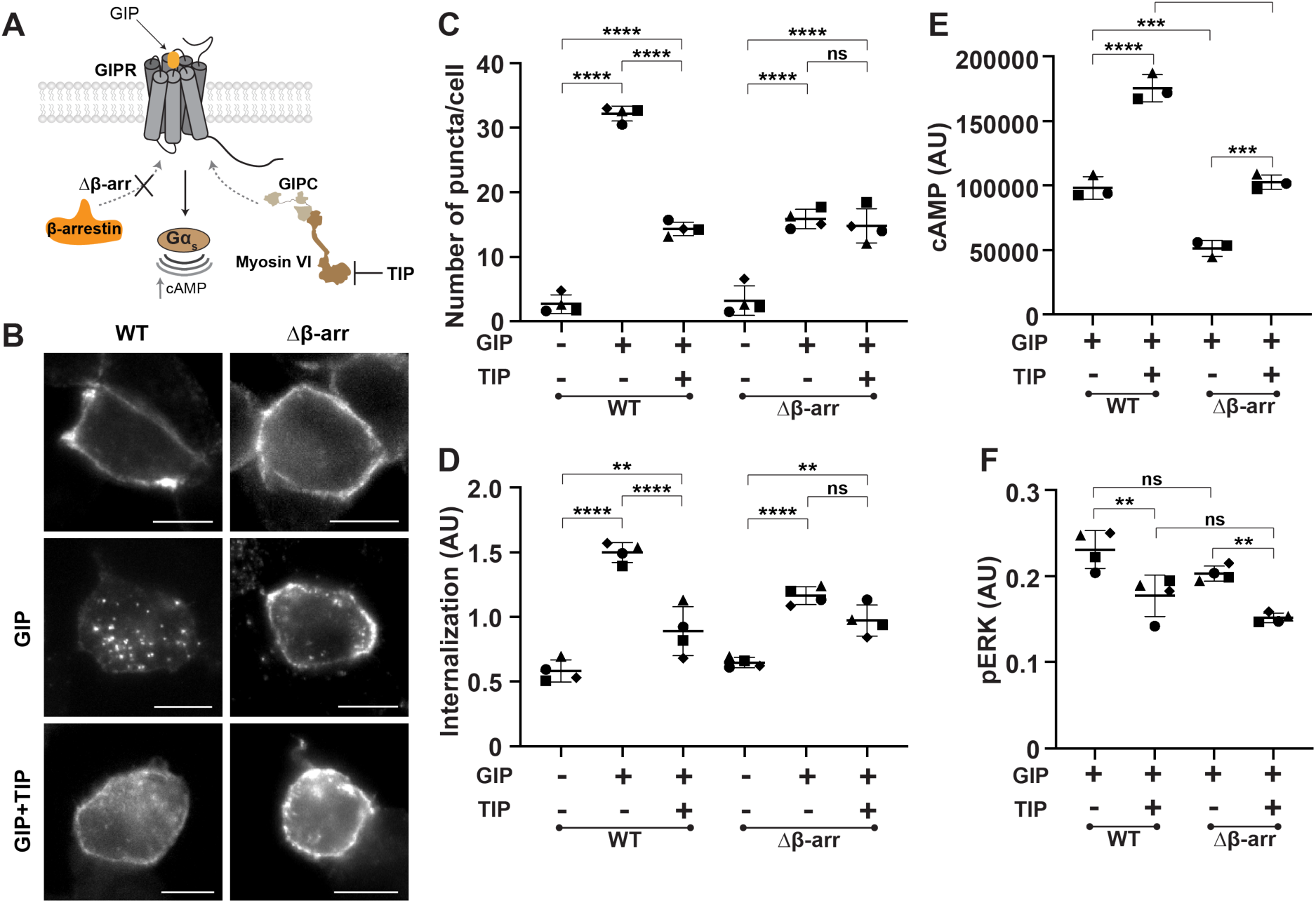
Myosin VI and β-arrestin synergistically regulate GIPR internalization and signaling. (A) Schematic representation of GIPR regulation by myosin VI and β-arrestin. GIPR C-tail has a PDZ binding motif (PBM) that binds PDZ adaptor GIPC, which in turn recruits cytoskeletal motor myosin VI. GIPR also recruits β-arrestin through multiple phosphorylation sites in its C-tail. Δβ-arr indicates β-arrestin 1/2 knockdown in HEK293 cells. TIP is a chemical inhibitor of myosin VI activity. (B) Internalization of GIPR with N-terminal FLAG tag, visualized using anti-FLAG antibody and secondary AlexaFlour 546. HEK293 wild-type (WT) and Δβ-arr cells expressing GIPR were stimulated with GIP (1 µM) for 15 min. Cells were treated with TIP (100 µM) to block myosin VI activity. Scale bar, 10 µm. (C-D) Number of puncta/cell (C) and cytosol-to-cell surface intensity (D) for GIPR internalization in HEK293 WT or Δβ-arr cells under basal, GIP or GIP+TIP conditions. Data is represented as mean +/-SD for four biological replicates. One-way ANOVA with Tukey’s post hoc test, ****p < 0.0001, **p < 0.01. (E-F) cAMP levels (E) and pERK levels (F) in HEK293 WT or Δβ-arr expressing GIPR and stimulated with GIP +/-TIP for 15 min. Mean +/-SD for three biological replicates. One-way ANOVA with Tukey’s post hoc test, ***p < 0.001, **p < 0.01, *p < 0.05.

### GIPC is recruited to GIPR prior to agonist treatment

- We have previously shown that a PDZ adaptor protein, GIPC, recruits myosin VI to PBMs in GPCR C-tails (20). Hence, we examined GIPC localization in cells expressing GIPR. GIPC-TagRFP fusions co-expressed with N-terminal FLAG-tagged GIPR in both WT and Δβ-arr HEK293 cells, show co-localization of Tag-RFP and Alexa Fluor 647 (M2 FLAG monoclonal + fluorescent secondary) at the cell plasma membrane prior to GIP treatment (Fig. 2A, Fig. S1A). Agonist (GIP; 1 μM; 15 min) stimulation results in receptor internalization, with persistent GIPC colocalization observed in endomembrane compartments in WT cells. While β-arrestin knockdown and TIP pretreatment both result in substantially lower internalization (Fig. 1B-D), GIPC colocalization persists both at the plasma membrane and endomembrane compartments. Dual inhibition of myosin VI and β-arr abolishes GIP-stimulated internalization, resulting in GIPC colocalizing with GIPR at the plasma membrane. Together, these data demonstrate that GIPC is recruited to the receptor independent of agonist, β-arrestin, or myosin VI activation. In contrast, β-arrestin does not localize to GIPR in the absence of GIP treatment (Fig. 2B, Fig. S1B). tagRFP-β-arrestin2 (β-arr2) fusions remain cytosolic when co-expressed with FLAG-tagged GIPR (visualized with M2 FLAG antibody mixed with Alexa Fluor 647 secondary). GIP stimulation results in β-arrestin recruitment to GIPR endosomes, regardless of TIP pretreatment or knockdown of endogenous β-arrestin. GIP stimulation with dual inhibition of myosin VI and β-arrestin mimics unstimulated cells, with cytosolic β-arrestin and receptor at the plasma membrane. We conclude that consistent with its established mode of interaction (26), β-arrestin is only recruited to the receptor following GIP stimulation, with visible colocalization with GIPR at endosomal compartments.

**Figure 2:**
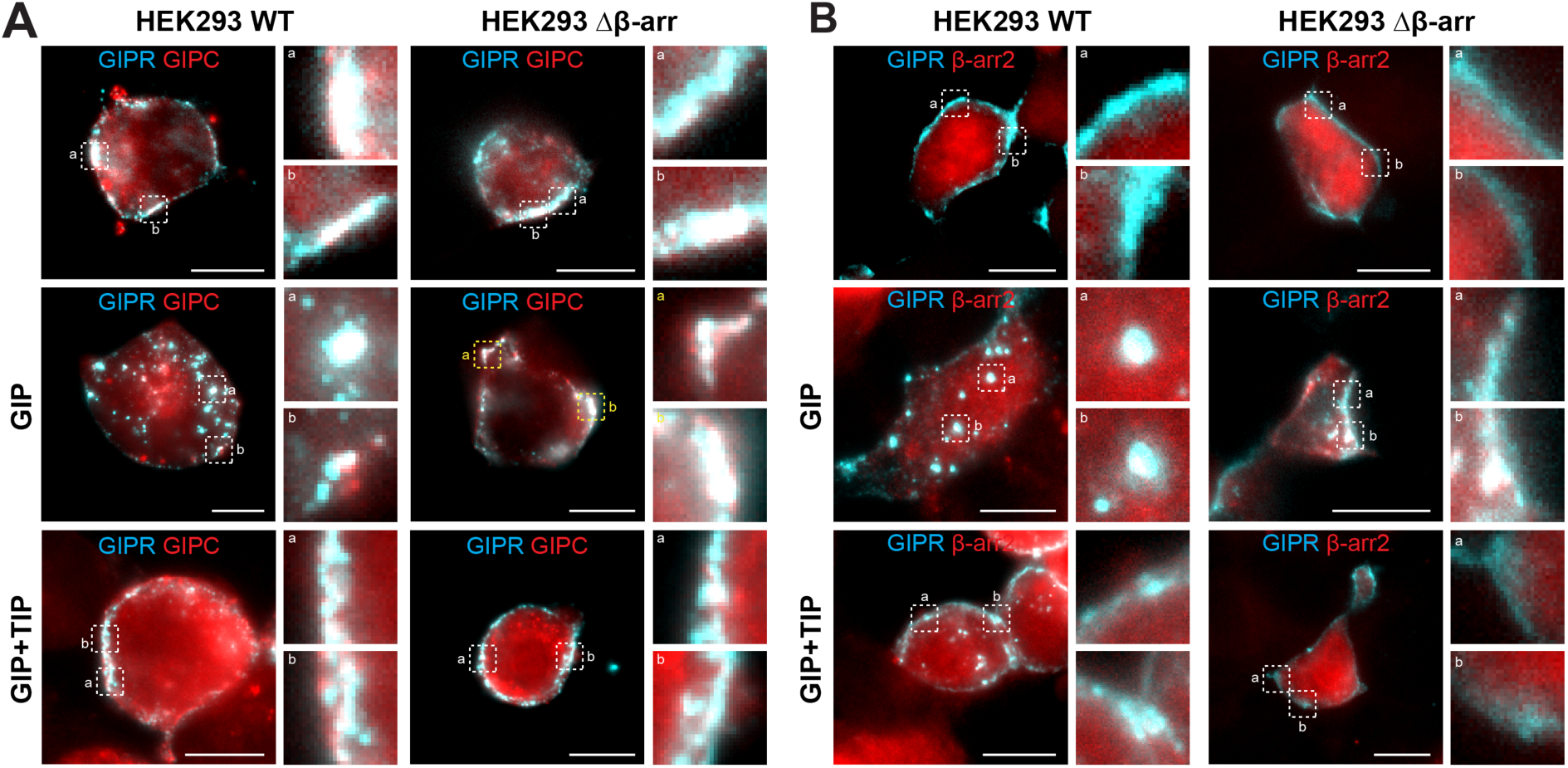
GIPC recruitment to GIPR is independent of β-arr. Merged images for colocalization of GIPR with GIPC (A) or GIPR with β-arr2 (B) in HEK293 WT or Δβ-arr cells, under basal condition or GIP stimulation +/-TIP for 15 min. HEK293 cells expressing FLAG-GIPR with GIPC or β-arr2 were visualized by immunostaining the receptor with anti-FLAG antibody and secondary AlexaFlour 647. Insets display magnified view of the areas marked with dashed squares. Scale bar, 10 μm. Refer to Fig. S1 for images of the individual channels.

### β-arrestin and GIPC interactions with GIPR are both essential for receptor internalization

**-** Given the colocalization of GIPC and β-arrestin with GIPR on endosomes, we examined the significance of their interactions with the GIPR C-tail for receptor internalization. The GIPC PBM site was abolished by truncating the last five amino acids (ΔPBM - LESYC), whereas β-arrestin engagement was limited by phospho-null mutations (S>A) (Fig. 3A). Removing the PBM motif substantially reduced GIPR internalization (Fig. 3B), with reduced punctae per cell (Fig. 3C) and cytosol-to-membrane fluorescence ratio (Fig. 3D). The phosphonull mutant also demonstrated lower GIPR internalization (Fig. 3B-D), albeit to a lesser extent than the PBM truncation. We also observe similar level of internalization for the phosphomimetic variant of GIPR, compared to the wild-type receptor (Fig. S2). These data further substantiate the dual role of β-arrestin and GIPC interactions in GIPR internalization.

**Figure 3:**
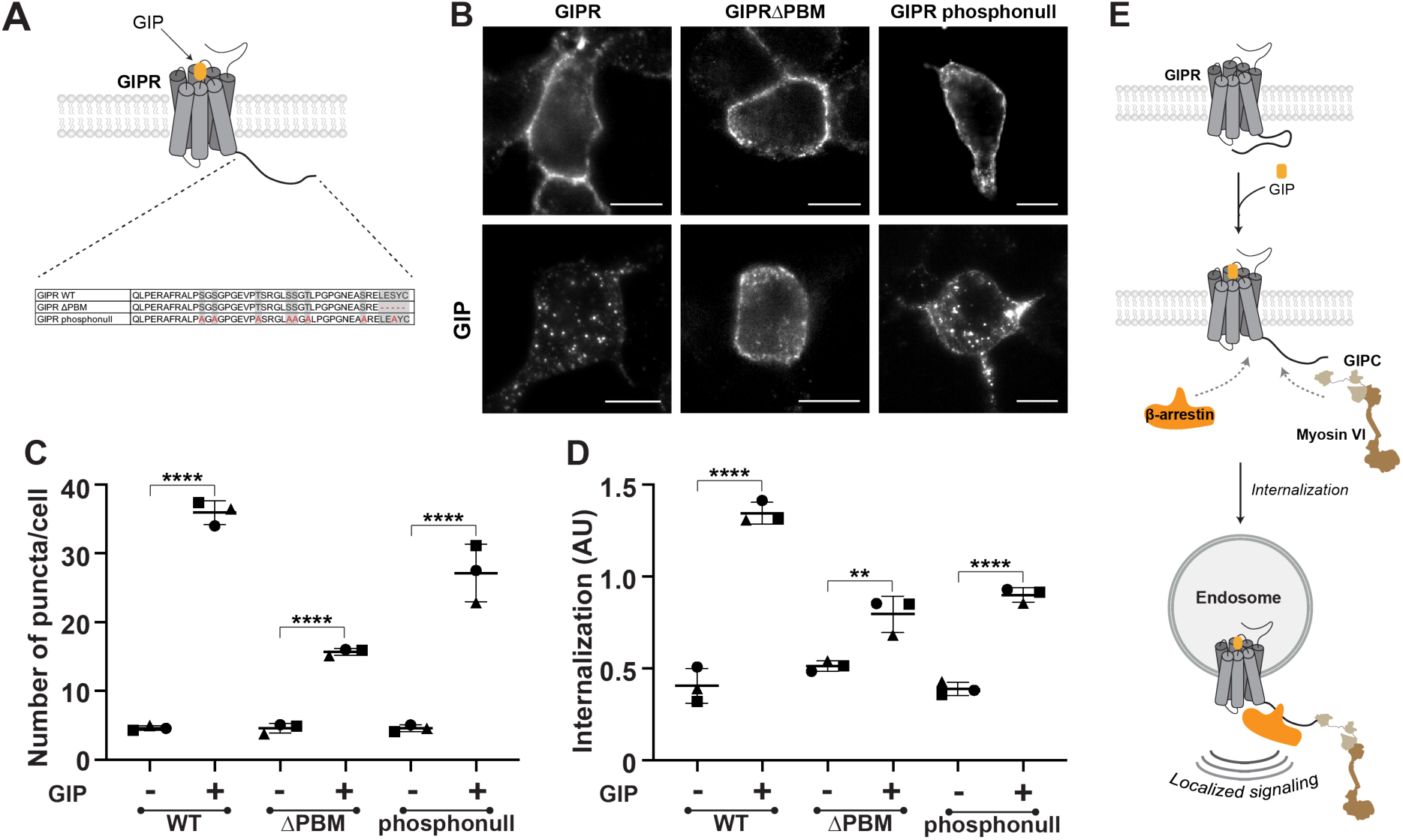
GIPR internalization is dependent on PBM sequence and phosphorylation in the C-tail. (A) Schematic representation of GIPR variants used in internalization assay. GIPR ΔPBM was created by removing the last five residues of the C-tail. Phosphorylation sites highlighted in gray were mutated to alanine to generate GIPR phosphonull construct. (B) Representative images of HEK293 WT cells expressing GIPR WT, ΔPBM and phosphonull, under basal and GIP stimulation. Refer to Fig. S3 for GIPR phospomimetic data. For surface labeling of the receptor, cells were incubated with anti-FLAG antibody and AlexaFlour 546/647. Scale bar, 10 μm. (C-D) Number of puncta/cell (C) and ratio for cytosol-to-cell membrane intensity (D) for GIPR WT, ΔPBM, phosphonull under basal and GIP-stimulated cells. Data is represented as mean +/-SD for three biological replicates. One-way ANOVA with Tukey’s post hoc test. ****p < 0.0001, **p < 0.01. (E) Model schematic for regulation of GIPR internalization and signaling by synergistic engagement of GIPC-myosin VI and β-arrestin.

### GIPC engagement of GIPR C-tail is enhanced by β-arrestin

- We used an *in vitro* FRET-based assay to further examine interactions between GIPC, β-arrestin and GIPR C-tail (Fig. 4A). Recombinant mCerulean-tagged GIPC (donor) demonstrates significant FRET (quantified using FRET ratio) with mCitrine-tagged GIPR C-tail (Fig. 4B). Both phosphomimetic (pm) and phosphonull (pn) versions of the recombinant GIPR C-tail show similar FRET ratios, supporting GIPC interactions with GIPR C-tail prior to agonist stimulation (Fig S3B). Addition of recombinant β-arrestin enhances this interaction, suggesting that GIPC and β-arrestin cooperatively bind the GIPR C-tail. To examine the functional consequences of this cooperativity, we used an established *in vitro* motility assay that reports on myosin VI activity as witnessed by the speed of actin filaments gliding on a myosin coated surface (Fig. 4C). As previously reported, GIPC stimulates actin gliding speeds (20, 27), with an increase observed upon addition of a phosphomimetic GIPR C-tail peptide (Fig. 4D, Fig. S3C). Addition of recombinant β-arrestin further augments actin gliding speed (Fig. 4D). Together, these data demonstrate synergies between GIPC and β-arrestin in stimulating myosin VI activity through engagement of the GIPR C-tail.

**Figure 4:**
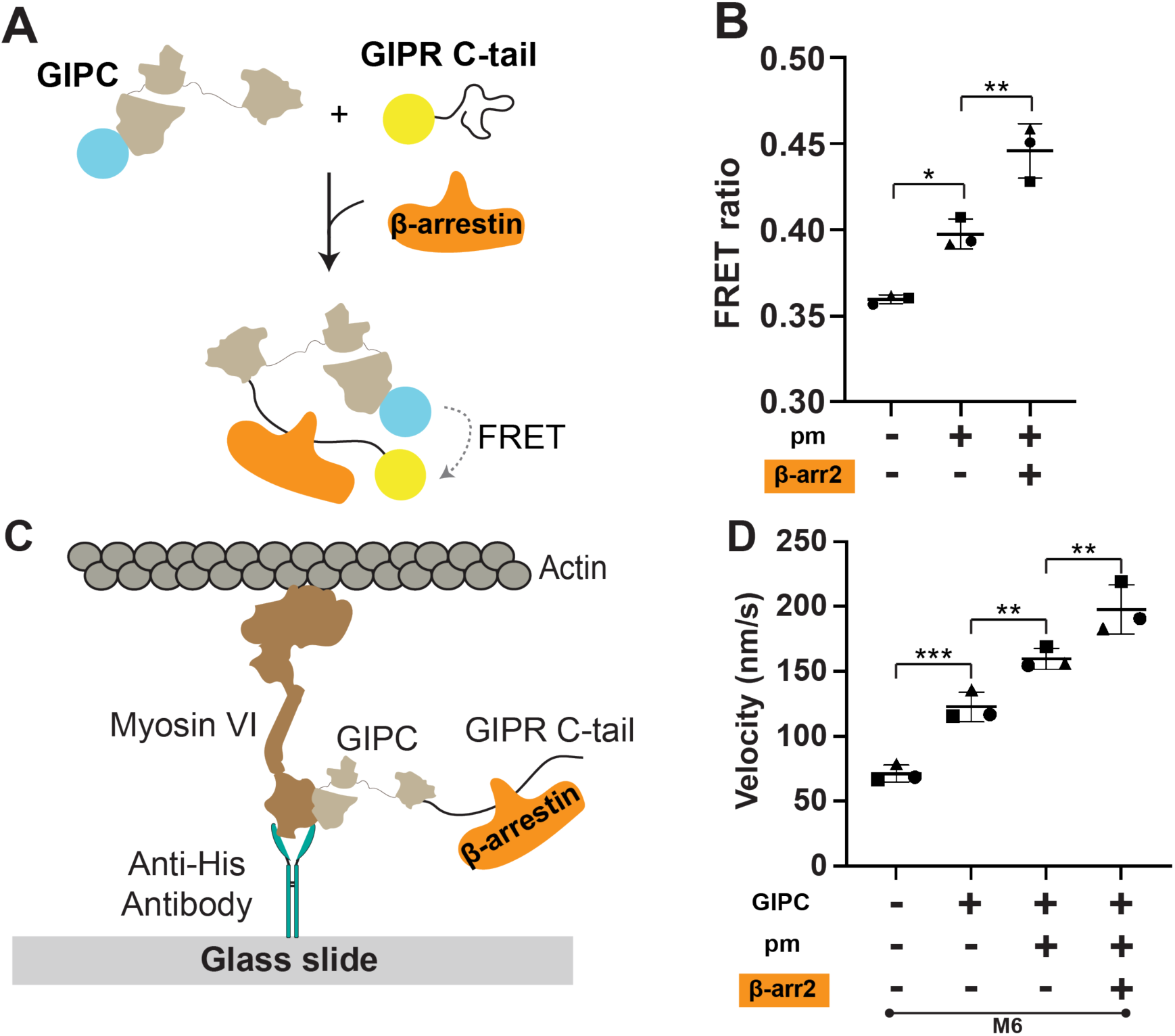
β-arrestin enhances GIPC engagement of the GIPR C-tail and myosin VI activity. (A) Schematic of bimolecular FRET experiment between mCer-GIPC and mCit-GIPR C-tail. Wild-type (wt), phoshonull (pn), and phoshphomimetic (pm) versions of the GIPR C-tail were used (Fig. S3A). Purified β-arrestin (β-arr2) was included with GIPR pm condition. (B) FRET ratio (525/475 nm) for bimolecular FRET between mCer-GIPC (30 nM) and mCit-GIPR C-tail pm (300 nM) +/- β-arr2 (300 nM). Data is represented as mean +/- SD for three biological replicates. One-way ANOVA with Tukey’s post hoc test. **p < 0.01, *p < 0.05. (C) Schematic for surface motility assay to determine myosin VI velocity. (D) Myosin VI speed with GIPC (2 μM), GIPR C-tail pm (100 μM) and β-arr2 (1 μM). Data is represented as mean +/- SD for three biological replicates. One-way ANOVA with Tukey’s post hoc test. ***p < 0.001, **p < 0.01. Refer to Fig. S3 for FRET and motility data of GIPR C-tail WT and pn.

### Myosin VI mediates insulin secretion at GIPR

- Previous studies have shown that β-arrestin is necessary for enhanced insulin release in response to GIP stimulation in pancreatic islets (26). Our data show that β-arrestin augments GIPC recruitment to the GIPR C-tail upstream of myosin VI activation. Hence, we tested whether myosin VI activity plays a role in GIP stimulated insulin secretion using a rat insulinoma cell line (INS-1 832/3) (Fig. 5A). TIP treatment (10 μM) enhanced GIP stimulated (1 μM) cAMP release (Fig. 5B) and insulin secretion (Fig. 5C) supporting a role for myosin VI in desensitization of physiological responses of GIPR.

**Figure 5:**
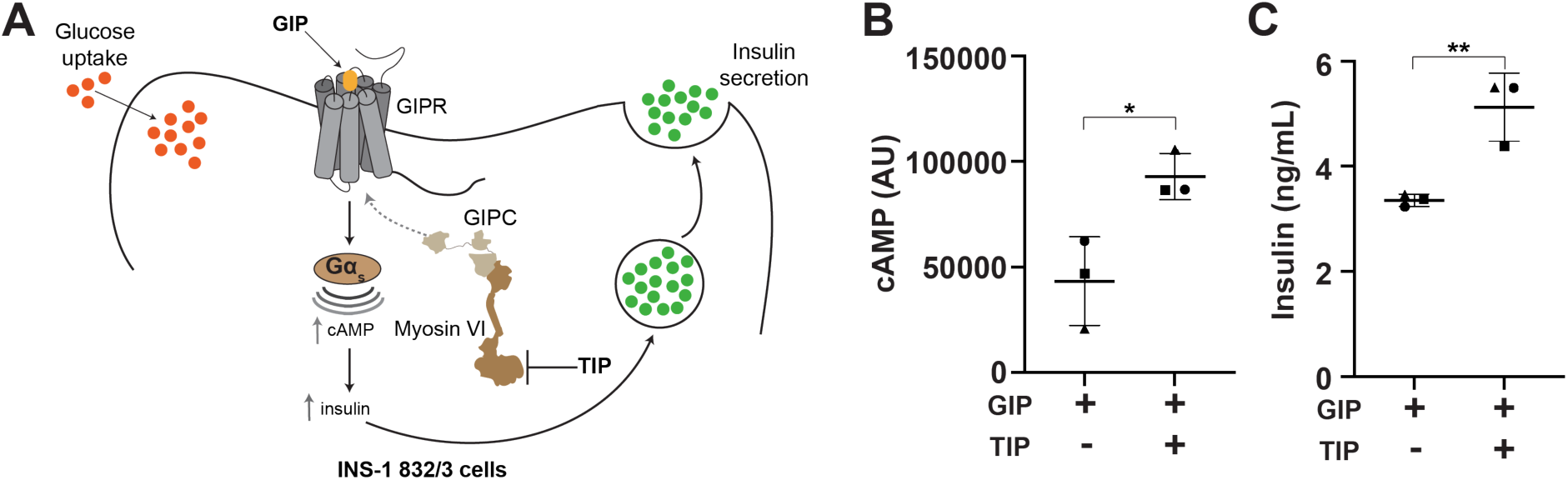
Myosin VI desensitizes incretin signaling at GIPR. (A) Schematic depiction of glucose-stimulated insulin secretion at GIPR. In presence of glucose, GIP stimulation of GIPR leads to cAMP production, resulting in insulin secretion from pancreatic beta cells. (B) cAMP response in INS cells, under GIP stimulation +/- TIP for 15 min. (C) Glucose-dependent insulin secretion by INS cells, in response to GIP stimulation +/- TIP for 30 min. Mean +/- SD for three biological replicates. Unpaired t-test. **p < 0.01, *p < 0.05.

## Discussion

GIPR signaling is essential for glucose-stimulated insulin release from pancreatic beta cells, making it a prominent target in metabolic disorders. Insulin response to the native agonist, GIP is influenced by the spatiotemporal trafficking of this receptor. Unlike prototypical GPCRs (3), β-arrestin knockdown has previously been shown to result in only modest effects on GIPR internalization (23). Here, we address this knowledge gap by identifying a novel incretin receptor trafficking mechanism through the engagement of the cytoskeletal motor, myosin VI. β-arrestin binds GIPR through established phosphorylation sites in the receptor C-tail (19). We find that β-arrestin binding augments GIPR C-tail interactions with a PDZ-adaptor GIPC (Fig. 4B). GIPC in turn recruits and activates the cytoskeletal motor myosin VI to promote GIPR endocytosis (Fig. 4D). Blocking myosin VI activity promotes GIPR retention at the plasma membrane with consequent enhanced cAMP signaling and reduced pERK1/2 activation from endosomal compartments (Fig. 1B-F). β-arrestin knockdown has been previously demonstrated to enhance GIP-stimulated insulin release in pancreatic islets (28). We find that myosin VI activity promotes desensitization of the insulin response in rat insulinoma cells, demonstrating the significance of this pathway in incretin receptor physiology (Fig. 5C).

Our study highlights the direct intersection between two trafficking pathways at the level of the receptor. We observe that GIPC, the adaptor for myosin VI, co-localizes with GIPR, irrespective of agonist stimulation (Fig. 2A). However, β-arrestin colocalization with GIPR is dependent on agonist treatment (Fig. 2B) (26). This suggests that although both pathways are important for GIPR internalization, GIPC-myosin VI engagement with GIPR likely occurs prior to β-arrestin recruitment. Nonetheless, binding of β-arrestin further strengthens the interactions between GIPC-myosin VI and GIPR, as evident in our results from binding and motility assays (Fig. 4B, Fig. 4D). Together, our findings demonstrate that these two trafficking pathways, rather than being mutually exclusive, operate synergistically to drive GIPR internalization.

The GPCR C-tail is an intrinsically disordered region (IDR) that autoregulates receptor signaling and engages effectors (β-arrestin/PDZ adaptors) to influence trafficking (30). We have limited structure-function insight into the C-tail of GPCRs, given the inherent conformational heterogeneity of IDRs. Their structural plasticity allows IDRs to engage with interacting partners through versatile and atypical mechanisms (31). Recent studies have highlighted the role of GPCR IDRs in the autoregulation of receptor activity by masking G protein access to the cytosolic cavity (32, 33). Heng et al. revealed that the β2AR C-tail interacts with intracellular loops (ICLs) to autoinhibit the receptor under basal conditions and blocking access to Gs (32). We have previously demonstrated that intramolecular interactions release spatial distance constraints in β2AR ICL3, facilitating Gs coupling (33). Further, biophysical studies demonstrate changes in GPCR C-tail conformation upon phosphorylation or effector binding, with unknown functional consequences (34, 35). Here, our data suggest cooperative engagement of β-arrestin and GIPC with the GIPR C-tail (Fig. 4B). We speculate that binding of one effector reduces conformational entropy of the C-tail, increasing access to another interacting partner.

GIPR represents a receptor that engages both β-arrestin and GIPC through cooperative interactions. Individually, GPCRs can differentially engage both PDZ-adpators and β-arrestin. Receptors either weakly (class A) or robustly (class B) recruit β-arrestin depending on the number and pattern of phosphorylation sites in the GPCR C-tail (36). Supporting the significance of C-tail phosphorylation and β-arrestin in receptor trafficking, our phosphonull version of GIPR results in lower internalization (Fig. 3B-D). In parallel, various GPCRs contain different types of PDZ binding motifs (I, II, and III) that we have demonstrated to bind with varying strengths to GIPC, in turn differentially stimulating myosin VI activity (8, 20). In this study, deletion of the PBM (GIPRΔPBM) site in GIPR leads to diminished receptor internalization (Fig. 3B-D). Supporting the significance of the GIPR PBM, this variant is associated with a metabolic phenotype (37). Together, we propose that the diversity of PBM and phosphomotifs enable differential regulation of GPCR trafficking to tune the spatiotemporal signaling response.

## Materials and Methods

### Cell Culture

HEK293T Flp-In T-Rex cells (referred to as HEK293 cells, ThermoFisher) and HEK293 β-arrestin 1/2 knockdown cells (referred to as HEK293 Δβ-arr) (22) were maintained in Dublecco’s modified eagle medium (DMEM, ThermoFisher), supplemented with GlutaMax (Thermo Fisher), 20 mM HEPES buffer (Corning) and 10% fetal bovine serum (FBS, GenClone). Cells were routinely cultured in 10 cm tissue-culture treated plates at 37°C with 5% CO_2_.

INS-1 832/3 (INS) cells (MilliporeSigma) were cultured in RPMI-1640 media, supplemented with 10% FBS, 10mM HEPES, 2 mM L-Glutamine, 1 mM sodium pyruvate and 0.05 mM β-mercaptoethanol. *Spodoptera frugiperda* insect cells (referred to as Sf9 cells, Thermo Fisher) were maintained at 28°C in Sf900-II media (ThermoFisher) supplemented with antibiotic-antimycotic (ThermoFisher).

### Plasmid construction

All plasmid constructs used in this study were generated using standard cloning procedures. Human GIPR, GIPC and β-arrestin2 (β-arr2) were cloned into the pcDNA5/FRT vector backbone. All GIPR constructs in pcDNA contained FLAG and mNeonGreen (mNG) tags at the N-terminus of the GPCR. Last five residues (L462-C466) were deleted for generating GIPRΔPBM. Phosphorylation sites in the C-tail were mutated to glutamine or alanine for phosphomimetic or phosphonull versions of the receptor, respectively (refer to Fig. 3A and Fig. S2A for the C-tail sequences). For the colocalization experiment, tagRFP was included at C-terminus of GIPC and at N-terminus of β-arr2.

Constructs used for bimolecular FRET and motility assays contained a FLAG tag for purification and were cloned into pBiex backbone. GIPR C-tail (Q422-C466) sequence with mCit tag at N-terminus was cloned into pBiex. Phosphonull (pn) and phosphomimetic (pm) variants of GIPR C-tail were cloned by mutating the phosphorylation residues to alanine or glutamine, respectively and contained mCit tag at the N-terminus. Full length M6 (M6-pBiex), GIPC-mCer-pBiex, and GIPC-pBiex were cloned as described previously (27).

### Internalization assay

HEK293 WT or Δβ-arr cells were seeded on poly-L-lysine coated glass coverslips in 35-mm plates (GenClone) at 30% density. The next day, cells were transfected with 4 μL polyethylenimine (PEI; PolySciences) and 1 μg of DNA of the indicated GIPR construct. For the co-localization experiment, 1 μg FLAG-mNG-GIPR and 0.5 μg of either GIPC-tagRFP or tagRFP-β-arr2 were transfected with 6 μL PEI. Internalization assay was carried out 24 hours post-transfection for all conditions, except GIPR + β-arr2 co-localization, transfection was allowed for 48 hours.

Cells were blocked in DMEM with 0.1% BSA for 15 min at 4°C prior to labeling. Anti-FLAG antibody from Millipore Sigma or GenScript performed similarly for labeling surface receptors. Anti-FLAG antibody was incubated with a secondary AlexaFlour 546 or 647 (Thermo Fisher) in DMEM with 0.1% BSA at 4°C for 60 min. Anti-FLAG+AlexaFlour complex was used to label surface receptors in HEK293 cells expressing GIPR at 4°C for 60 min. Next, cells were stimulated with GIP (1 μM; GenScript) for 15 min. When using TIP, cells were pretreated with TIP (100 μM; 15 min; MilliporeSigma) prior to agonist stimulation and TIP was included during GIP stimulation. Cells were fixed in 4% formaldehyde for 10 min at room temperature. Coverslips were then washed in PBS and mounted on glass slide using Prolong Diamond Anti-Fade (ThermoFisher). Coverslips were sealed the next day with valap (vaseline/lanolin/paraffin) and imaged on a Nikon Eclipse Ti inverted epifluorescence microscope with 100X oil immersion objective (1.4 numerical aperture) equipped with Evolve EMCCD camera (Photometrics) or PCO.edge sCMOS camera using Nikon Elements software (Nikon). Z-stacks (0.6 μm step size) were acquired and maximum intensity projection (MIP) were created using Fiji (ImageJ)(38). Number of particles and cytosol-to-surface intensity was determined using Fiji, as described previously(20). At least three biological replicates were carried out for each condition and 10-20 cells were analyzed per each condition for every biological replicate.

### cAMP assay

HEK293 WT or Δβ-arr cells were seeded in 6-well plate at 30% cell density and transfected with 1 μg FLAG-mNG-GIPR using 4 μL PEI for 24 hrs. For cAMP in INS cells, untransfected cells were seeded in a 6-well plate and processed the next day. On the day of the assay, cells were harvested in culturing media and centrifuged at 350 g for 5 min at room temperature. Next, cells were resuspended in stimulation buffer (PBS, 0.2% glucose, 0.5 mM ascorbic acid), cell count was determined using a hemocytometer (Countess II) and added to an opaque 384-well plate (Greiner Bio-One) at 1.5*10^4 cells/well. The samples were then processed according to the manufacturer’s instructions for the cAMP glo assay kit (Promega). Cells were stimulated with 1 μM GIP +/-10 μM TIP for 15 min at room temperature and lysis buffer (Promega) was added to stop the reaction. Absorbance was recorded on a plate reader (Tecan Spark) with 500 ms integration. Basal cAMP (no agonist) was subtracted from respective samples to calculate the amount of cAMP. Surface receptor expression levels in HEK293 WT or Δβ-arr were measured by recording fluorescence spectra on FluoroMax-4 spectrofluorometer (Horiba Scientific - excitation 470 nm, emission 495-600 nm, 4 nm bandpass). At least four technical replicates and three biological replicates were performed for each condition.

### phosho-ERK1/2 (pERK) assay

HEK293 WT or Δβ-arr cells were seeded in 35-mm plates at 30% confluency. The next day, media was exchanged to serum-free DMEM and cells were transfected with GIPR (1 μg) using 4 μL PEI. Cells were harvested in serum-free DMEM 24 hours after transfection and serum starvation. Cell density was determined using a hemocytometer (Countess II). Cells were then added to a white 384-well plate (Greiner Bio-One) at 1*10^4 cells/well. The pERK assay was carried out following the manufacturer’s instructions(Revvity). To stimulate, cells were incubated with 1 μM GIP for 15 min. To block myosin VI activity, TIP was added at 10 μM along with the agonist GIP. Reactions were quenched by adding lysis buffer (Revvity) and the plate was incubated at room temperature for 30 min with shaking at 500 rpm. Pre-mixed antibody solution was added to each well and incubated at room temperature for 16-20 hours. pERK levels were measured using HTRF-compatible plate reader (Molecular Devices - excitation 314 nm, cutoff 630 nm; emission 665 nm and 620 nm, cutoff 570 nm). pERK amount was calculated as the ratio between 665 nm and 620 nm values. At least four technical replicates and three biological replicates were included for each condition.

### Glucose-stimulated insulin secretion (GSIS) assay

INS cells were seeded in 12-well plates at 1*10^6 cells/well density and processed for the GSIS assay at ∼90% confluency. Cells were washed in Krebs-Ringer bicarbonate buffer (KRB buffer; 130 mM NaCl, 5 mM KCl, 1.2 mM CaCl_2_, 1.2 mM MgCl_2_, 1.2 mM KH_2_PO_4_, 25 mM NaHCO_3_, 20 mM HEPES, 0.2% BSA), followed by glucose starvation in KRB buffer with 2.5 mM glucose for two hours at 37°C. Next, cells were stimulated with 11 mM glucose +/-GIP (1 μM) for 30 min at 37°C. TIP was added at 10 μM to block myosin VI activity. Supernatant containing secreted insulin was collected and added to a 384-well plate. Insulin levels were measured according to the manufacturer’s instructions (Revvity) using an HTRF-compatible plate reader (Molecular Devices - excitation 314 nm, cutoff 630 nm; emission 665 nm and 620 nm, cutoff 570 nm). Briefly, antibody was added to the wells and incubated overnight before acquisition on the plate reader. Insulin concentration was calculated from an insulin standard curve. Buffer +/- TIP was used to subtract basal insulin secretion from respective test conditions. At least four technical replicates and three biological replicates were included for every condition.

### Bimolecular FRET

Sf9 cells at 2*10^6 cells/mL density were transfected with DNA for 7 μg GIPC-mCer, 5-7 μg mCit-GIPR C-tail (WT, pm or pn) or 10 μg β-arr2 and Escort IV transfection reagent (MilliporeSigma). Proteins were purified from Sf9 cells 72-hours post-transfection, as described previously(20). Briefly, cells were harvested and lysed in lysis buffer (20 mM imidazole pH 7.5, 7% sucrose, 200 mM NaCl, 4 mM MgCl_2_, 0.5 mM EDTA, 1 mM EGTA, 5 mM DTT, 0.5% IGEPAL, 5 μg/ml aprotinin, 5 μg/ml leupeptin, 5 μg/ml PMSF). The lysate was clarified by centrifugation at 176,000 g for 25 min at 4°C. Next, the supernatant was incubated anti-FLAG affinity resin (MilliporeSigma) for 60 min at 4°C. Then, the resin was washed three times in wash buffer (20 mM imidazole pH 7.5, 7% sucrose, 150 mM KCl, 5 mM MgCl_2_, 1 mM EDTA, 1 mM EGTA, 5 mM DTT, 5 μg/ml aprotinin, 5 μg/ml leupeptin, 5 μg/ml PMSF). Elution of the bound protein was carried out by incubating the resin with FLAG peptide (GenScript) overnight at 4°C. Verification and quantification of protein expression was done by running the purified protein samples on SDS-PAGE gel. FRET assays were performed with proteins diluted in FRET buffer (20mM HEPES pH 7.4, 25 mM KCl, 5 mM MgCl_2_). GIPC-mCer was used at 30 nM and mCit-GPCR C-tails and β-arr2 were added at 300 nM. WT, pm or pn variants of the GIPR C-tail were used where indicated. Spectra for FRET samples were recorded on a FluoroMax-4 spectrofluorometer (Horiba Scientific) by exciting at 430 nm (8 nm bandpass) and emission from 450 nm to 650 nm (4 nm bandpass and 1 nm intervals). mCit-GPCR C-tail spectra +/- β-arr2 were subtracted to account for cross-excitation. FRET ratio (525 nm/475 nm) was determined using peak intensity values for mCitrine (525 nm) and mCerulean (475 nm).

### Motility assay

Sf9 cells were transfected with DNA (7 μg myosin VI, 7 μg GIPC or 10 μg β-arr2) using Escort IV transfection reagent. Proteins were purified as described above. Purified β-arr2 was further desalted with a Zeba spin column (ThermoFisher), following the manufacturer’s protocol. GPCR C-tails (WT, pm or pn) were commercially synthesized (GenScript). Myosin VI motility assays were carried out as previously described(20). Briefly, flow chambers were created with collodion-coated coverslips attached to glass slides with strips of double-sided tape. Anti-His antibody (Qiagen, 0.02 mg/mL) in assay buffer (AB; 20 mM imidazole pH 7.5, 25 mM KCl, 4 mM MgCl_2_, 1 mM EGTA) was passed through the flow chamber and allowed to incubated for 4 min, followed by three washes with 10 μL AB-BSA (AB with 1 mg/mL BSA) and blocking with 10 μL AB-BSA for 2 min. Next, pre-mixed myosin VI (200 nM), GIPC (2 μM) and GIPR C-tail (100 μM) in AB-BSA were flown through the chamber and incubated for 4 min. For β-arr2 condition, β-arr2 (1 μM) in AB-BSA was flown separately through the chamber and incubated for 2 min. AB-BSA (10 μL) was used between each step to wash off unbound protein. Finally, actin motility mix (F-actin labeled with Alexa-647 phalloidin (ThermoFisher), 45 μg/ml catalase (EMD Millipore), 2 mM ATP, 1 mM phospho-creatine (MilliporeSigma), 0.1 mg/ml creatine phosphokinase (EMD Millipore), 0.6% glucose, 25 μg/ml glucose oxidase (EMD Millipore)), supplemented with GIPC and GIPR C-tail was flown through the flow chamber. A Nikon Eclipse Ti inverted epifluorescence microscope equipped with an oil immersion objective (100x, 1.4 NA) was used to record motility movies at 1 frame/sec for 3 min with PCO.edge sCMOS camera using Nikon Elements software (Nikon). Motility movies were analyzed using the MTrack plugin on Fiji. For each condition, 25-30 tracks were analyzed and all conditions were performed in three biological replicates.

### Statistical analysis

GraphPad Prism (version 10) was used to generate graphs and compute statistical significance. All data is represented as mean +/- SD and details of the statistical tests performed are mentioned in the figure legends.

## Acknowledgements

We thank Michael Ritt for his feedback on the manuscript. This research was funded by the NIH (R35-GM126940 to S.S. and 5K12GM119955-07 to N.M.P.).

## Supplementary Figure legends

**Figure S1:**
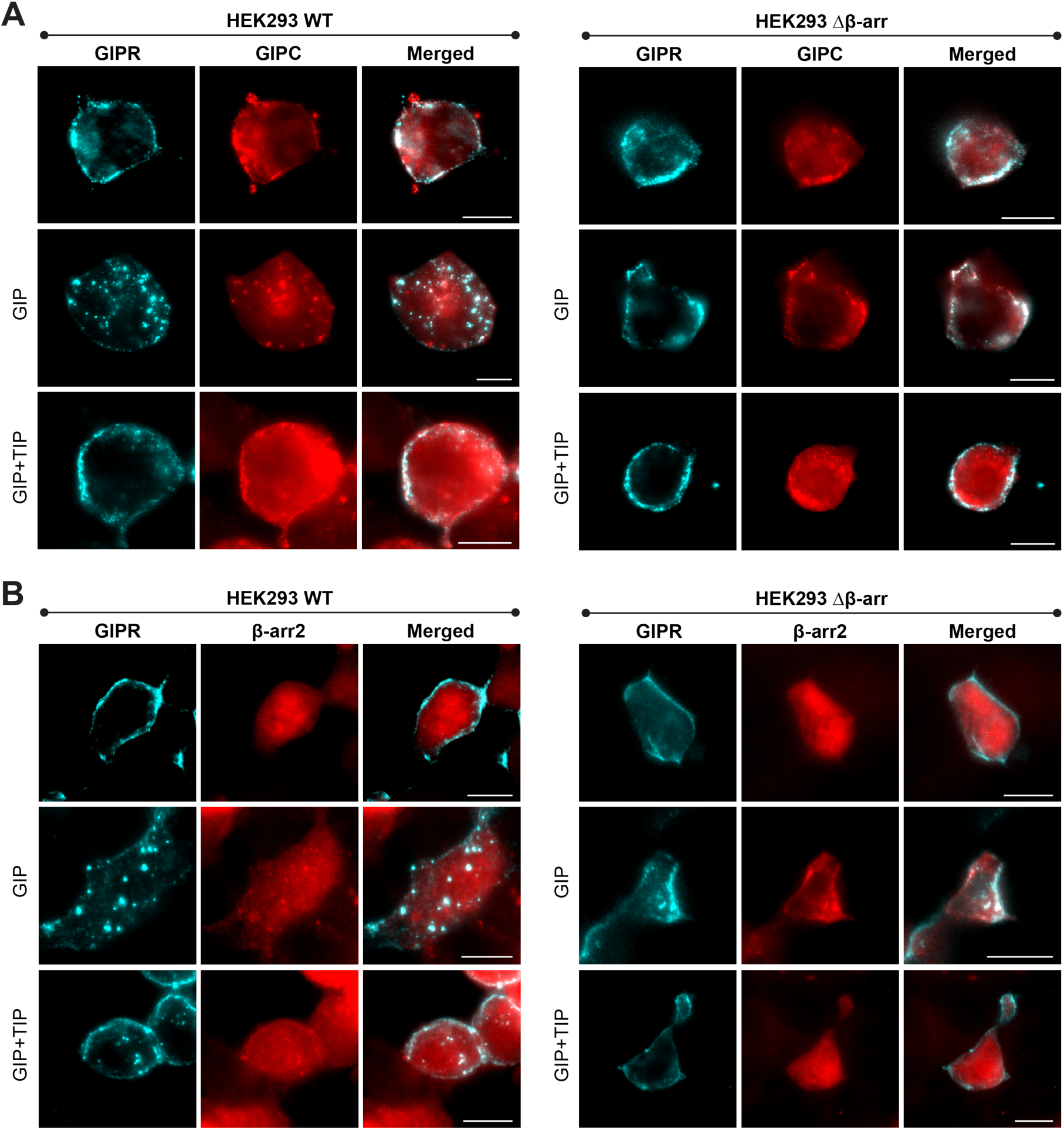
Colocalization of GIPC or β-arr2 with GIPR. HEK293 WT or Δβ-arr cells expressing FLAG-GIPR with GIPC-tagRFP (A) or tagRFP-β-arr2 (B) were visualized by immunostaining the receptor with anti-FLAG antibody + secondary AlexaFlour 647. Images are shown for basal or GIP stimulation +/- TIP conditions. Scale bar, 10 μm.

**Figure S2:**
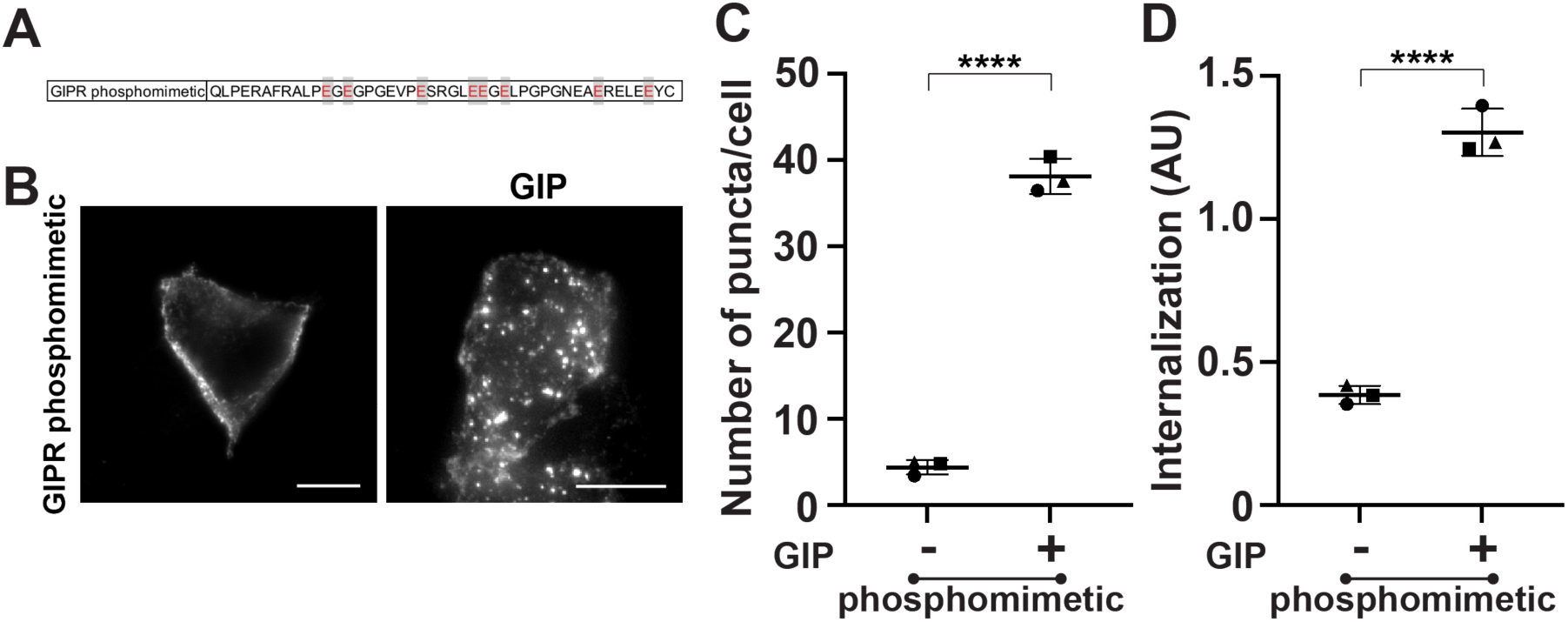
GIPR phosphomimetic undergoes complete internalization. (A) Schematic representation of GIPR phosphomimetic construct. Phosphorylation sites highlighted in gray were mutated to glutamine to generate phosphomimetic version of the receptor. (B) Representative images of HEK293 WT cells expressing GIPR phosphomimetic under basal and GIP stimulation. Cells were incubated with anti-FLAG antibody and AlexaFlour 546/647 for surface labeling of the receptor. Scale bar, 10 μm. (C-D) Number of puncta/cell (C) and ratio for cytosol-to-cell membrane intensity (D) for GIPR phosphomimetic under basal and GIP-stimulated cells. Data is represented as mean +/- SD for three biological replicates. One-way ANOVA with Tukey’s post hoc test. ****p < 0.0001.

**Figure S3:**
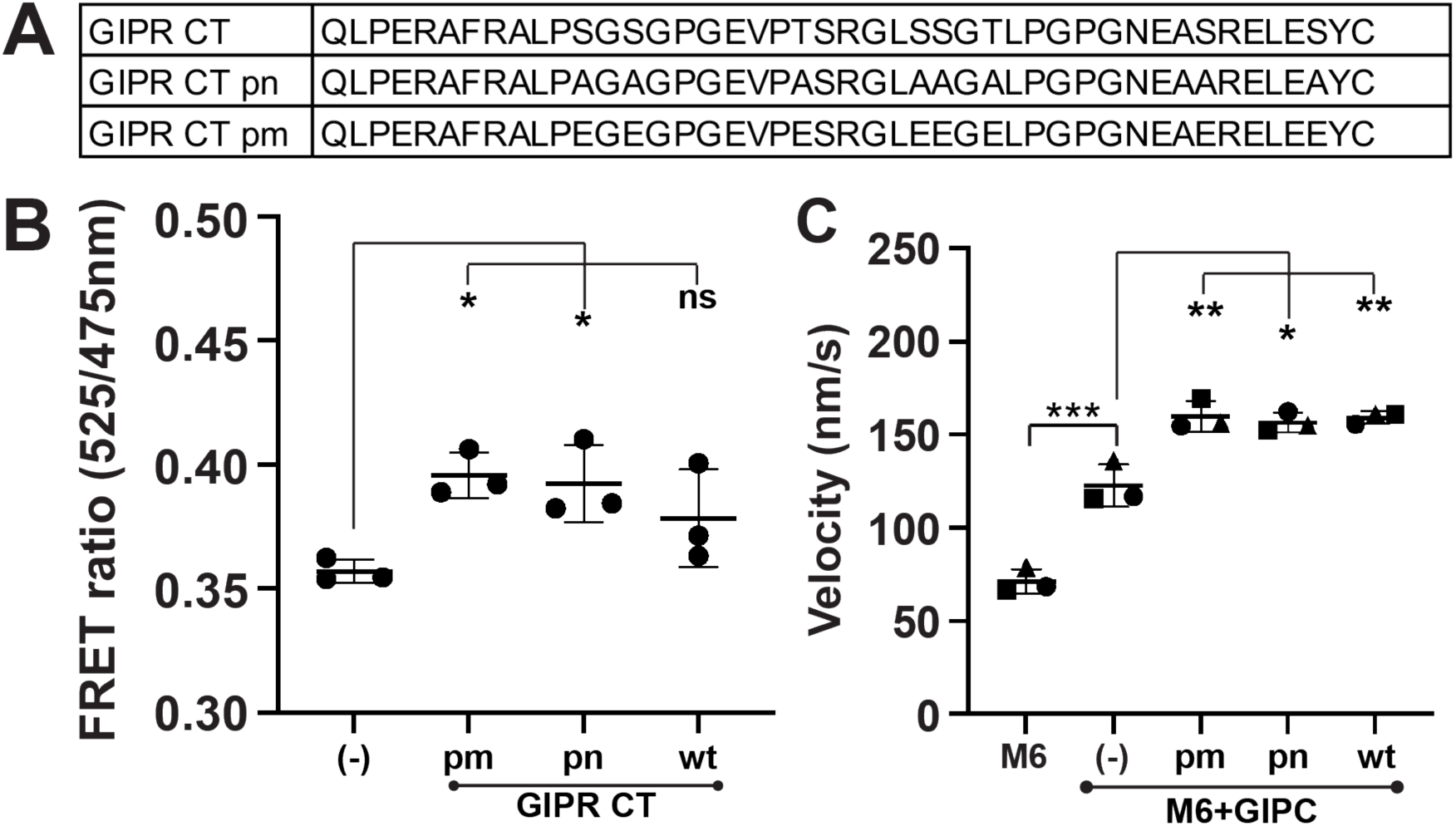
GIPR C-tail interacts with GIPC and activates myosin VI. (A) Sequence of GIPR CT (wild-type (wt), phoshonull (pn), and phoshphomimetic (pm)) used in the FRET and motility assays. (B) FRET ratio (525/475 nm) for bimolecular FRET between mCer-GIPC (30 nM) and mCit-GIPR C-tail (300 nM). Data is represented as mean +/-SD for three biological replicates. One-way ANOVA with Dunnett’s post hoc test. *p < 0.05. (C) Myosin VI motility with GIPC and GIPR C-tail (wt, pn, or pm). Data is represented as mean +/-SD for three biological replicates. One-way ANOVA with Tukey’s post hoc test. ***p < 0.001, **p < 0.01, *p < 0.05.

